# Functions of Gle1 are governed by two distinct modes of self-association

**DOI:** 10.1101/2020.08.20.259564

**Authors:** Aaron C. Mason, Susan R. Wente

**Author notes:** Corresponding author: Susan R. Wente.

## Abstract

Gle1 is a conserved, essential regulator of DEAD-box RNA helicases, with critical roles defined in mRNA export, translation initiation, translation termination, and stress granule formation. Mechanisms that specify which, where and when DDXs are targeted by Gle1 are critical to understand. In addition to roles for stress-induced phosphorylation and inositol hexakisphosphate (IP_6_) binding in specifying Gle1 function, Gle1 oligomerizes via its N-terminal domain in a phosphorylation-dependent manner. However, a thorough analysis of the role for Gle1 self-association is lacking. Here, we find that Gle1 self-association is driven by two distinct regions: a coiled-coil domain and a novel ten amino acid aggregation prone region, both of which are necessary for proper Gle1 oligomerization. By exogenous expression in HeLa cells, we tested the function of a series of mutations that impact the oligomerization domains of the Gle1A and Gle1B isoforms. Gle1 oligomerization is necessary for many, but not all aspects of Gle1A and Gle1B function, and the requirements for each interaction domain differ. Whereas the coiled-coil domain and aggregation prone region additively contribute to competent mRNA export and stress granule formation, both self-association domains are independently required for regulation of translation under cellular stress. In contrast, Gle1 self-association is dispensable for phosphorylation and non-stressed translation initiation. Collectively, we reveal self-association functions as an additional mode of Gle1 regulation to ensure proper mRNA export and translation. This work also provides further insight into the mechanisms underlying human *gle1* disease mutants found in prenatally lethal forms of arthrogryposis.

## Introduction

Throughout the gene expression pathway, the fate of a protein-coding messenger (m)RNA transcript is determined by interactions between its specific complement of RNA-binding proteins and other cellular machinery. During this process, step-wise changes to messenger ribonucleoprotein (mRNP) complexes are achieved through the remodeling actions of RNA-dependent DEAD-box ATPases (termed Dbps in *Saccharomyces cerevisiae*, DDXs in humans) (1). Providing an additional layer of regulation, some Dbps/DDXs require specific cofactors that modulate their ATPase activities (2). Among the known cofactors, the essential conserved multidomain protein Gle1 is a uniquely versatile Dbp/DDX modulator. Gle1 is involved in facilitating proper nuclear mRNA export, translation, and the stress response (3–10).

In human cells, at least two different Gle1 isoforms, A and B, result from alternative splicing of a single *GLE1* pre-mRNA transcript (11). Differing exclusively in their C-terminal regions, only Gle1B contains a 39 amino acid extension that interacts with Nup42 on the cytoplasmic face of the nuclear pore complex (NPC) (12–15). Through the interaction with Nup42 and other NPC-associated factors, Gle1B shows steady state localization to the nuclear rim, whereas Gle1A is predominantly cytoplasmic and only at the nuclear rim when Gle1B is absent (3, 11). Through differing expression levels, protein interactions and subcellular localization, Gle1A and Gle1B execute unique functions. Gle1B facilitates the terminal step of mRNA export through the NPC by activating DDX19B in an inositol hexakisphosphate (IP_6_)-dependent manner (4, 9, 10, 13). In the cytoplasm, Gle1A is required for proper stress granule (SG) formation and translation initiation during cellular stress through IP_6_-independent modulation of DDX3 (3, 5, 6, 16). In *S. cerevisiae*, IP_6_-dependent Gle1 regulation of Dbp5, the orthologue of DDX19B, also plays a role during translation termination (6). However, a translation termination function for human Gle1A or Gle1B has not been reported. Substantial Gle1 structural and functional analysis maps the residues responsible for Dbp5/DDX19B and IP_6_ interactions to the C-terminal region, with no contribution of its N-terminal domains to ATPase stimulation (4, 7, 9, 12). Likewise, activation of DDX3 ATPase activity *in vitro* also requires only the Gle1 C-terminal domain (16).

Dbps/DDXs and their specific cofactors are also dynamically regulated by their oligomerization state, which directly alters their intrinsic activity and also provides new binding surfaces for other protein interactions. DDX3, and its *S. cerevisiae* orthologue Ded1, functions as a trimer to unwind RNA duplexes (17, 18). Ded1 oligomerization is disrupted by eIF4G and the Ded1/eIF4G complex also acts as a functional helicase but with reduced activity as compared to the trimeric Ded1 (18). DDX3 interaction with Ezrin inhibits its helicase activity and increases its ATPase activity (19), with Ezrin’s oligomeric state being influenced by phosphorylation (20). As further examples across a range of species, the cold shock RNA helicase, CsdA, from *E. coli* requires its C-terminal dimerization domain for activity and stability (21). In *T. thermophilus*, HerA exists as a heptameric and a hexameric ring with binding of NurA and nucleotide promoting a hexameric oligomerization state. Additionally, NurA functions as a dimer that tightly binds to hexametric HerA and enhances the affinity of HerA-NurA for dsDNA (22, 23).

Although the C-terminal region of Gle1 is necessary and sufficient for its modulation of DDX activities, the Gle1 N-terminal region self-associates both *in vitro* and *in vivo*, and is required for proper Gle1 localization and function (7). Indeed, the Gle1 N-terminal region has multiple reported protein-protein interaction attributes, including a short 29 residue Nup155 binding motif (24), followed by a 123 residue intrinsically disordered region (IDR) of unknown function and structure (16) and a 208 residue predicted coiled-coil domain (25). Biochemically, large 15 – 60 nm discs form *in vitro* from purified recombinant Gle1^1-360^, which are perturbed by a *gle1-Fin*_*major*_ mutation that introduces a proline, phenylalanine, glutamine (PFQ) insertion into the predicted coiled-coil domain (7). In human cells, Gle1 dimerization is observable by FRET and requires the N-terminal region. Deletion of the N-terminal region abolishes Gle1 nuclear rim localization, and correspondingly, nucleocytoplasmic shuttling, mRNA export, and *in vitro* self-association are all perturbed by the *gle1-Fin*_*major*_ mutation. Demonstrating the critical nature of Gle1 oligomerization for proper mRNA export, the *gle1-Fin*_*major*_ mutation underlies the fatal neurodevelopmental disease lethal congenital contracture syndrome-1 (LCCS1) (26). While studies in *S. cerevisiae* suggest that Gle1’s roles in translation initiation and termination are not dependent on the coiled-coil domain, this has not been investigated in human cells. Moreover, human Gle1’s N-terminal region contains a much longer IDR and is the site of dynamic, stress-induced phosphorylation that alters the formation of Gle1 oligomers *in vitro* (16), suggesting that other regions of the N-terminus contribute to oligomerization. Overall, further studies are needed to obtain a detailed understanding of the properties governing Gle1 self-association and its effect on isoform-specific functions.

Here we investigate the biochemical properties of Gle1 oligomerization and test how disrupting proper Gle1 oligomerization influences Gle1 function in many aspects of RNA metabolism. We find that two distinct N-terminal regions contribute to the overall oligomerization state of Gle1. The coiled-coil domain self-associates in a parallel orientation but is not sufficient to generate high molecular weight oligomers. Conversely, an aggregation prone region in the unstructured IDR is necessary for high molecular weight oligomerization. Mutations engineered to result in specific amino acid changes disrupt these two oligomerization domains and distinct functions *in vivo*. We show that proper Gle1 oligomerization is essential for proper Gle1 subcellular localization and function in mRNA export and heat-shock stress response.

## Results

### Gle1 contains two distinct regions that govern oligomerization

Our prior studies show that perturbation of the predicted N-terminal coiled-coil domain by the LCCS1-linked PFQ insertion impedes gle1-Fin_major_ oligomerization and nucleocytoplasmic shuttling, as well as poly(A)+ RNA export (7). The N-terminal Gle1 region also contains a Nup155 binding domain, a predicted IDR and a recently reported stress-induced phosphorylation cluster (Fig. 1A) (16, 24). To investigate the mechanism of human Gle1 self-association, we employed a set of biochemical approaches. First, using MARCOIL (27) to more accurately delineate the boundaries of the predicted coiled-coil domain, we found a near 100% probability of coiled-coil formation spanning amino acids 152 to 360 in the Gle1 N-terminal region (Fig. 1B).

**Figure 1.**
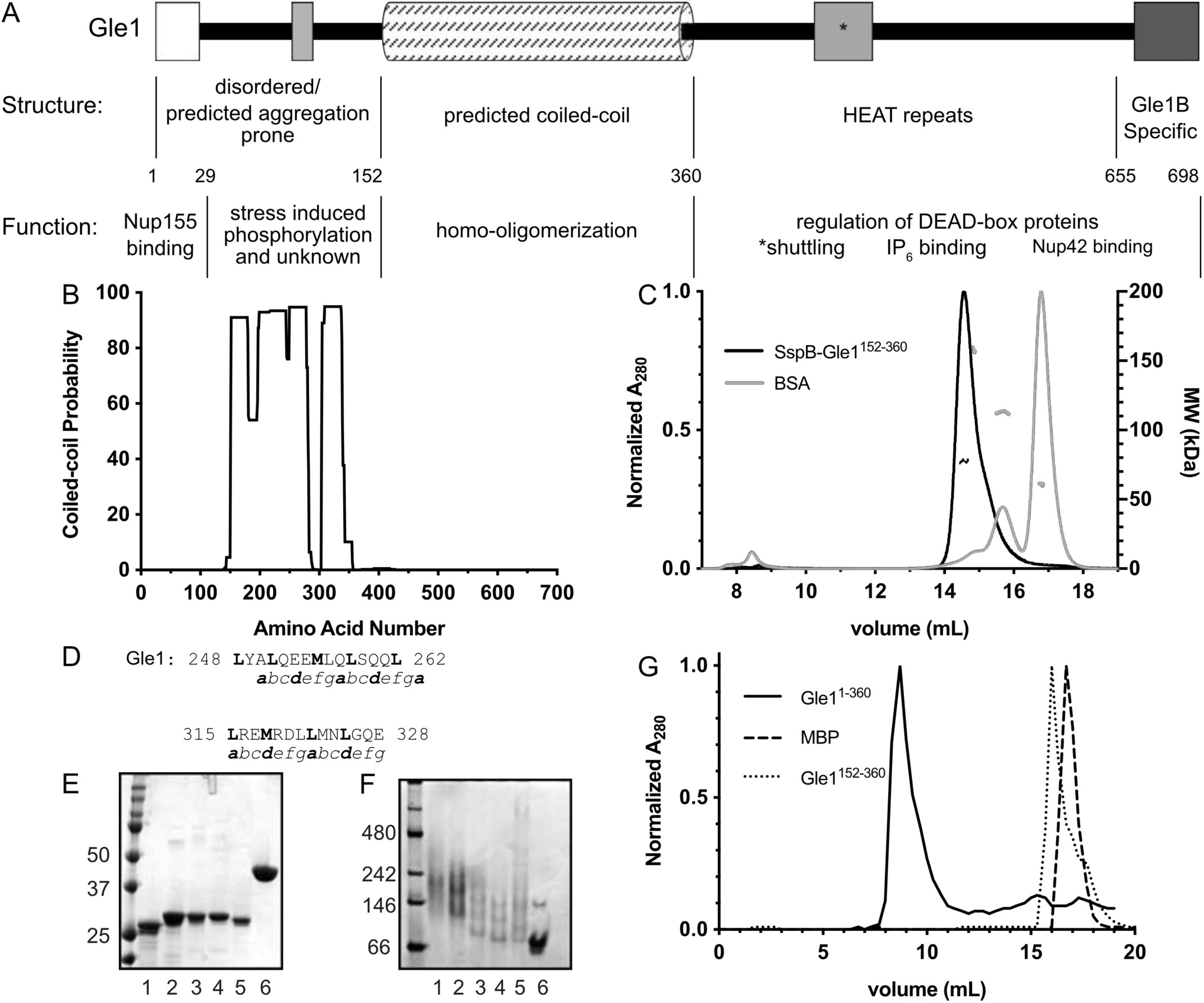
Coiled-coil domain contributes to Gle1 oligomerization. (A) Human Gle1 domain organization schematic. (B) MARCOIL predicted coiled-coil probability of Gle1 (C) Assessment tool for homodimeric Coiled-Coil ORientation Decision (ACCORD): Size-exclusion chromatography with multi-angle light scattering (SEC-MALS) of SspB-Gle1^152-360^ and BSA. SEC-MALS experiments were performed in buffer containing 20mM HEPES pH 7.6, 200mM NaCl, 5% glycerol, 0.5mM TCEP, 0.05% azide with a Superose6 10/300 GL column connected to a Wyatt Dawn8+ system. (D) Predicted putative heptad repeat of Gle1. (E, F) Putative heptad repeat mutants analyzed by SDS-PAGE (E) or Native PAGE (F). Lane 1: Gle1^152-360^, 2: Gle1^152-360 L248D L251D^, (2D) 3: Gle1 ^152-360 L248D L251D L262D L315D M318D^, (4D) 4: Gle1^152-360 L248D L251D L258D L262D L315D M318D^, (6D), Gle1^152-360 L248D L251D L258D L262D L315D M318D L322D L325D^ (8D), 6: MBP (G) Size-exclusion chromatography analysis of Gle1^1-360^, Gle1^152-360^, and MBP.

To date, no high-resolution structural studies are reported for the N-terminal Gle1 regions. Coiled-coil domains associate in either a parallel or anti-parallel orientation (28). To roughly discern how Gle1 domains are relatively positioned in a Gle1 oligomer, we probed the orientation required for Gle1 self-association by utilizing a technique termed Assessment tool for homodimeric Coiled-Coil ORientation Decision (ACCORD) (29). This method is based on forced dimerization by the *Escherichia coli* protein SspB when the respective coiled-coil domain being analyzed is expressed as a fusion protein with SspB. The resulting purified chimeric protein is then analyzed by SEC-MALS. Coiled-coil domains associating in a parallel orientation result in a single dimerization event, whereas anti-parallel orientation results in two dimerization interfaces that produce a tetramer or higher ordered complex. Analysis of the purified SspB-Gle1^152-360^ fusion protein by SEC-MALS revealed a sharp peak with a calculated molecular weight in agreement with a dimer (Fig. 1C). Therefore, we concluded that the Gle1 coiled-coil domain self-associates in a parallel orientation.

Coiled-coil domains are also predicted by a common heptad repeat of (*a-b-c-d-e-f-g*)_n_, wherein the *a* and *d* positions are predominantly hydrophobic residues that form the oligomer’s hydrophobic core (28). To identify the heptad repeats present in the Gle1 coiled-coil domain, the amino acid sequence from 1-360 was submitted to the MARCOIL web-based server for comparison with a database of known parallel-oriented coiled-coil interactions (27). MARCOIL confirmed the presence of the canonical heptad repeat in Gle1 (Fig. 1D), and predicted the interacting amino acids in the hydrophobic core of the interacting Gle1 coiled-coils. Based on those predictions, we mutated the coding sequence for a series of hydrophobic residues in the *a* and *d* positions in the context of Gle1^1-360^ to result in the introduction of aspartic acid (D) residues. The resulting purified proteins were analyzed by SDS-PAGE and Native PAGE (Fig. 1E,F respectively). As shown by native PAGE, successive introduction of charged amino acids into the predicted hydrophobic core resulted in faster electrophoretic migration, with the 8D protein fully collapsing the slower migrating bands that are indicative of Gle1 oligomers. Thus, Gle1 oligomerization *in vitro* required the interaction of parallel-oriented coiled-coil domains.

Analysis of recombinant Gle1^1-360^ by SEC and negative stain EM shows it elutes in the void volume as large, higher ordered oligomeric discs of 15 to 60 nm diameter (7). To test whether the coiled-coil domain alone is sufficient to form these large discs, we compared the SEC profiles of Gle1^152-360^ and Gle1^1-360^. As shown in Figure 1G, Gle1^1-360^ eluted in the void volume, consistent with previous reports. However, Gle1^152-360^ did not elute in the void volume, but rather eluted as a sharp peak in later fractions (Fig. 1G). This suggested that the coiled-coil domain alone is not sufficient to form high molecular weight oligomers.

Since expression of Gle1^152-360^, containing only the coiled-coil domain, did not produce high molecular weight oligomers, we speculated that a different region in the N-terminal region of Gle1 mediates the disc formation. The IDR in the N-terminal region was analyzed for molecular recognition features (DICHOT; (30)), short linear peptide motifs (ELM Prediction; (31)), prion-like domains (PrionW; (32)), and aggregation hotspots (AGGRESCAN; (33)). AGGRESCAN predicted a potential aggregation hot spot from amino acids 45-54 in the unstructured IDR between the Nup155 binding domain and the coiled-coil domain (Fig. 2A). To assess the contribution of this 10 amino acid stretch to the formation of higher ordered oligomers, this region was deleted from Gle1^1-360^ and the resulting recombinant purified protein was analyzed by SEC. Gle1^1-360 Δ45-54^ generated an SEC elution profile with two peaks split between the void and later fractions corresponding to the approximate molecular weight of monomeric Gle1^1-360 Δ45-54^, as compared to MBP (Fig. 2B). The protein in the later peak was further analyzed by circular dichroism (CD) to determine if the 10 amino acid deletion altered structural conformation or stability. Gle1^1-360^ and Gle1^1-360 Δ45-54^ produced similar CD spectra that are suggestive of a strong alpha helical protein (Fig. 2C). This data was consistent with a fraction of Gle1^1-360Δ45-54^ eluting in the later peak due to disruption of high ordered oligomerization, and not as the result of overall protein unfolding.

**Figure 2.**
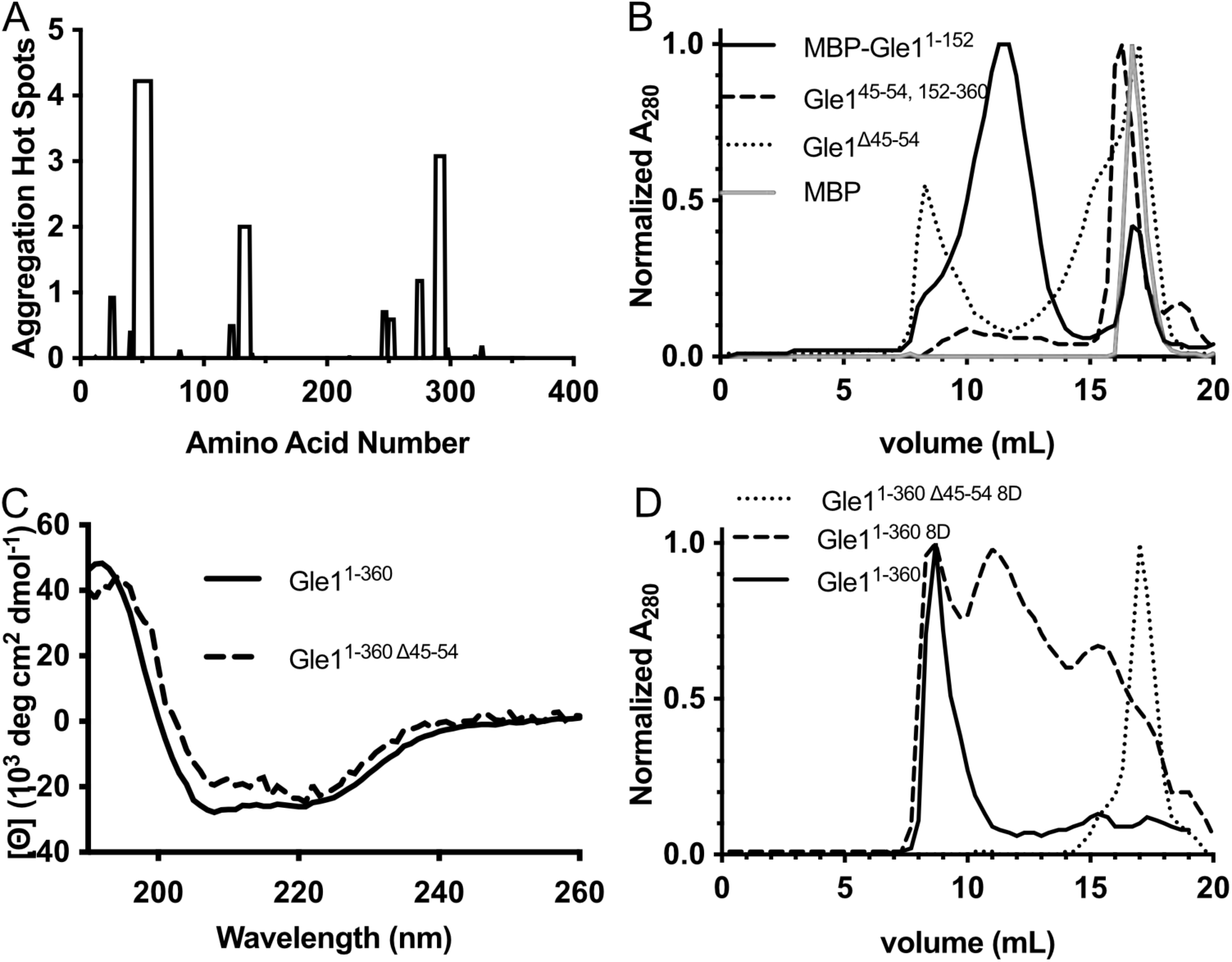
Characterization of an aggregation prone region of Gle1. (A) Results of Gle1 sequence submitted to AGGRESCAN server reveals a potential aggregation prone region. (B) Size-exclusion chromatography analysis of MBP-Gle1^1-152^, Gle1^45-^54, 152^-360^, Gle1^1-360 45-54^, and MBP. (C) Circular dichroism spectra of Gle1^1-360^ and the non-void volume peak of Gle1^1-360 45-54^. (D) Disrupting both the coiled-coil domain and aggregation prone region of Gle1 impairs oligomerization. Size-exclusion chromatography analysis of Gle1^1-360^, Gle1^1-360 45-54^, Gle1^1-3608D^.

The oligomerization potential of the N-terminal domain excluding the coiled-coil domain, but containing the aggregation hot spot and complete IDR, was also analyzed by SEC. As the resulting Gle1^1-152^ protein was unstable by itself, purification was aided by the addition of an N-terminal MBP tag. Surprisingly, the SEC elution profile of MBP-Gle1^1-152^ showed a single peak with a retention time close to the void volume, suggesting higher ordered oligomers (Fig. 2B). To examine if amino acids 45-54 were alone sufficient to drive the higher ordered oligomers, amino acids 45-54 were expressed in frame with the coiled-coil domain (Gle1^45-^54, 152^-360^). Analysis of Gle1^45-^54, 152^-360^ by SEC revealed an elution profile similar to that of Gle1^152-360^, with no indication of higher order oligomerization (Fig. 2B). These results demonstrated that the aggregation prone region of amino acids 45-54 is necessary but not sufficient in combination with the coiled coil domain to drive higher ordered Gle1 oligomerization. Rather, additional minor aggregation prone regions within amino acids 1-152 are likely required to promote formation of the large, higher order oligomers that elute in the void volume of wild type Gle1^1-360^ (Fig. 2A).

Based on this analysis, the coiled-coil domain and the aggregation prone region in the N-terminal region of Gle1 both contribute to Gle1 oligomerization. To investigate the effect of disrupting the coiled-coil domain within the context of the complete N-terminal region, the 8D mutations described in Fig. 1D were placed in the context of Gle1^1-360^ and analyzed by SEC. The SEC elution profile of Gle1^1-360 8D^ revealed a broad elution profile with multiple peaks including a peak in the void volume (Fig. 2D). This result indicated that the coiled-coil domain, while not sufficient to produce oligomers alone, contributes to higher ordered Gle1 oligomerization. From comparing the elution profile of Gle1^1-360 8D^ and that of Gle1^1-360 Δ45-54^ (which suggested that oligomerization was marginally destabilized by loss of the aggregation hot spot), we speculated that further combining both the deletion of the aggregation prone region and the 8D mutations would result in a Gle1 protein that is unable to oligomerize. Recombinant purified Gle1^1-360 Δ45-54 8D^ was therefore examined by SEC. As shown in Figure 2D, the elution profile of Gle1^1- 360 Δ45-54 8D^ showed a single peak late in the elution, indicative of monomerization. Therefore, *in vitro* oligomerization of Gle1 was dependent on both the N-terminal aggregation prone region and coiled- coil domain, and alteration of either domain impeded oligomerization.

### Gle1 oligomerization is required for proper subcellular localization

To examine the effects of disrupting Gle1 oligomerization *in vivo*, we first investigated the localization of the aggregation prone deletion and coiled-coil disrupting mutants in context of either Gle1A or Gle1B (Fig. S2). In HeLa cells, live cell imaging of exogenously-expressed GFP-Gle1A in the presence of endogenous Gle1 (both Gle1A and Gle1B) showed steady state localization in the cytoplasm without observable rim localization, as previously shown (Fig. 3A) (11). As expected, we found that exogenously-expressed GFP-gle1A-8D, GFP-gle1A-Δ45-54, and GFP-gle1A-Δ45-54-8D mutants also exhibited steady state localization in the cytoplasm (Fig. 3A, B). In the presence of endogenous Gle1, exogenously-expressed GFP-Gle1B was distributed pancellularly at steady state with marked nuclear rim localization (Fig. 3A), consistent with previous studies (11). Unexpectedly, GFP-gle1B-8D, GFP-gle1B-Δ45-54, and GFP-gle1B-Δ45-54-8D all showed steady state localization in the cytoplasm with a marked decrease in the nucleoplasm as compared to GFP-Gle1B (Fig. 3A, B).

**Figure 3.**
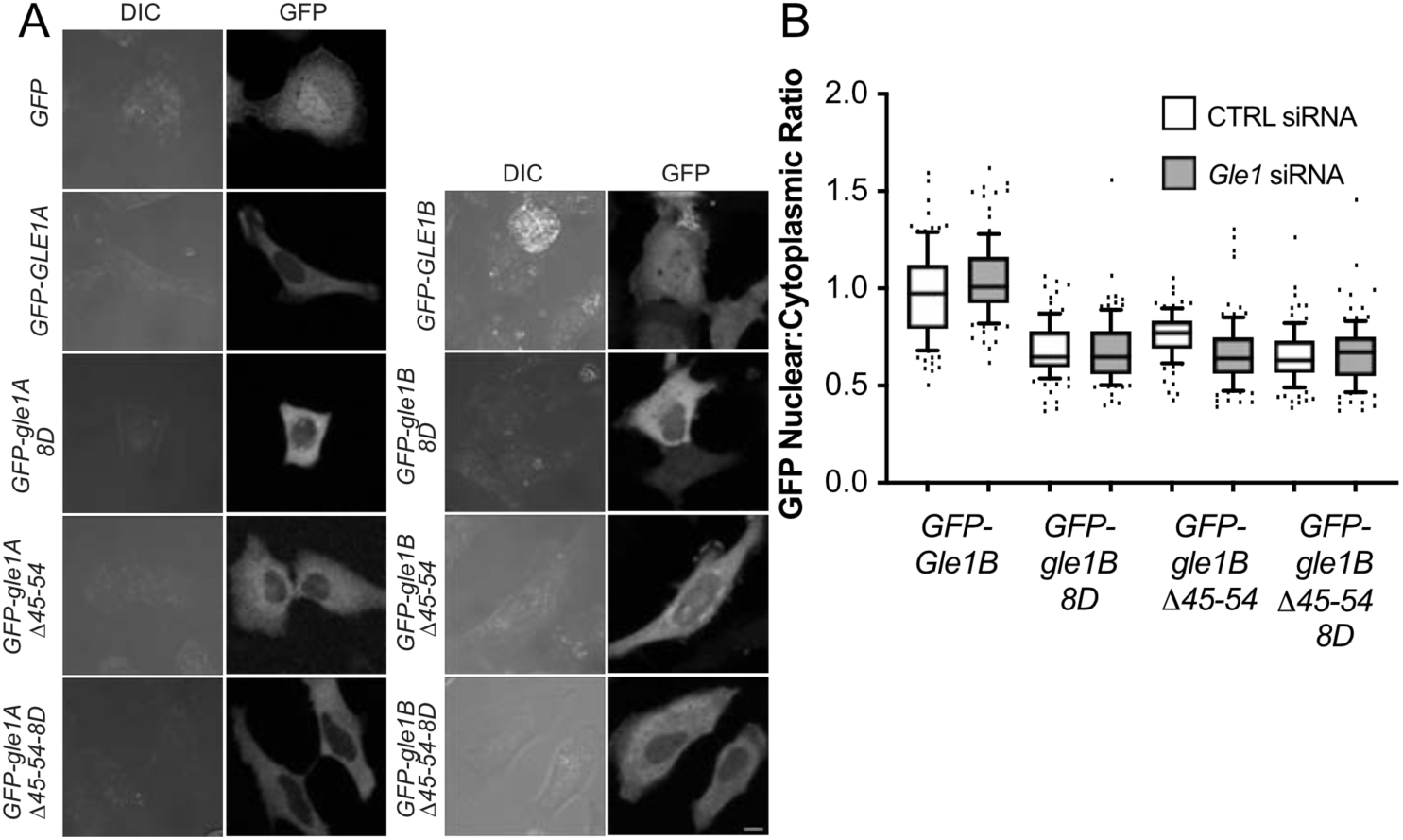
Gle1 oligomerization required for proper subcellular localization (A) HeLa cells transfected with *GFP, GFP-GLE1A, GFP-gle1A-8D, GFP-gle1-ΔA45-54, GFP-gle1A-Δ45-54-8D, GFP-GLE1B, GFP-gle1B-8D, GFP-gle1B-Δ45-54*, or *GFP-gle1B-Δ45-54-8D* were visualized by direct fluorescent live cell imaging. Scale bar: 10 µm. (B) Quantification of the GFP nuclear to cytoplasmic ratio of HeLa cells expressing *GFP-GLE1B, GFP-gle1B-8D, GFP-gle1BΔ-45-54, or GFP-gle1B-Δ45-54-8D* either CTRL or GLE1 siRNA treated.

Our previous studies show that the en masse deletion of Gle1B amino acids 40-400 perturbs localization to the nuclear rim, whereas the gle1-Fin_major_ variant that disrupts the coiled coil domain is still detected at the nuclear rim (7). To better define the oligomerization requirements for Gle1 nuclear rim localization, we conducted a detailed analysis of colocalization at the NPC by immunofluorescence with mAb414, which recognizes the FXFG repeat in a family of NPC proteins. First, Gle1A and its oligomerization-deficient variants were examined. In the presence of endogenous Gle1A and Gle1B, as noted, exogenously-expressed GFP-Gle1A does not compete for localization at the NPC (11). Consistent with those observations, exogenously-expressed GFP-Gle1A did not appear to colocalize with mAb414 in the presence of endogenous Gle1A and Gle1B, nor did any of the GFP-gle1A oligomerization variants. This was confirmed by fluorescence signal quantification, showing the peak GFP intensities clearly shifted toward the cytoplasmic side of the histogram relative to mAb414 (Fig. 3C). However, as reported, if *GLE1* is knocked down resulting in depletion of endogenous Gle1A and Gle1B, exogenously-expressed GFP-Gle1A localized to the nuclear rim (Fig. 4A) (3). In sharp contrast, following depletion of endogenous Gle1A and Gle1B, exogenously-expressed GFP-gle1A-8D, GFP-gle1A-Δ45-54, and GFP-gle1A-Δ45-54-8D were notably absent from the NPC, as demonstrated by their lack of colocalization with mAb414 (Fig. 4A). This data indicated that in the absence of Gle1B, Gle1A localization at the NPC requires oligomerization.

**Figure 4.**
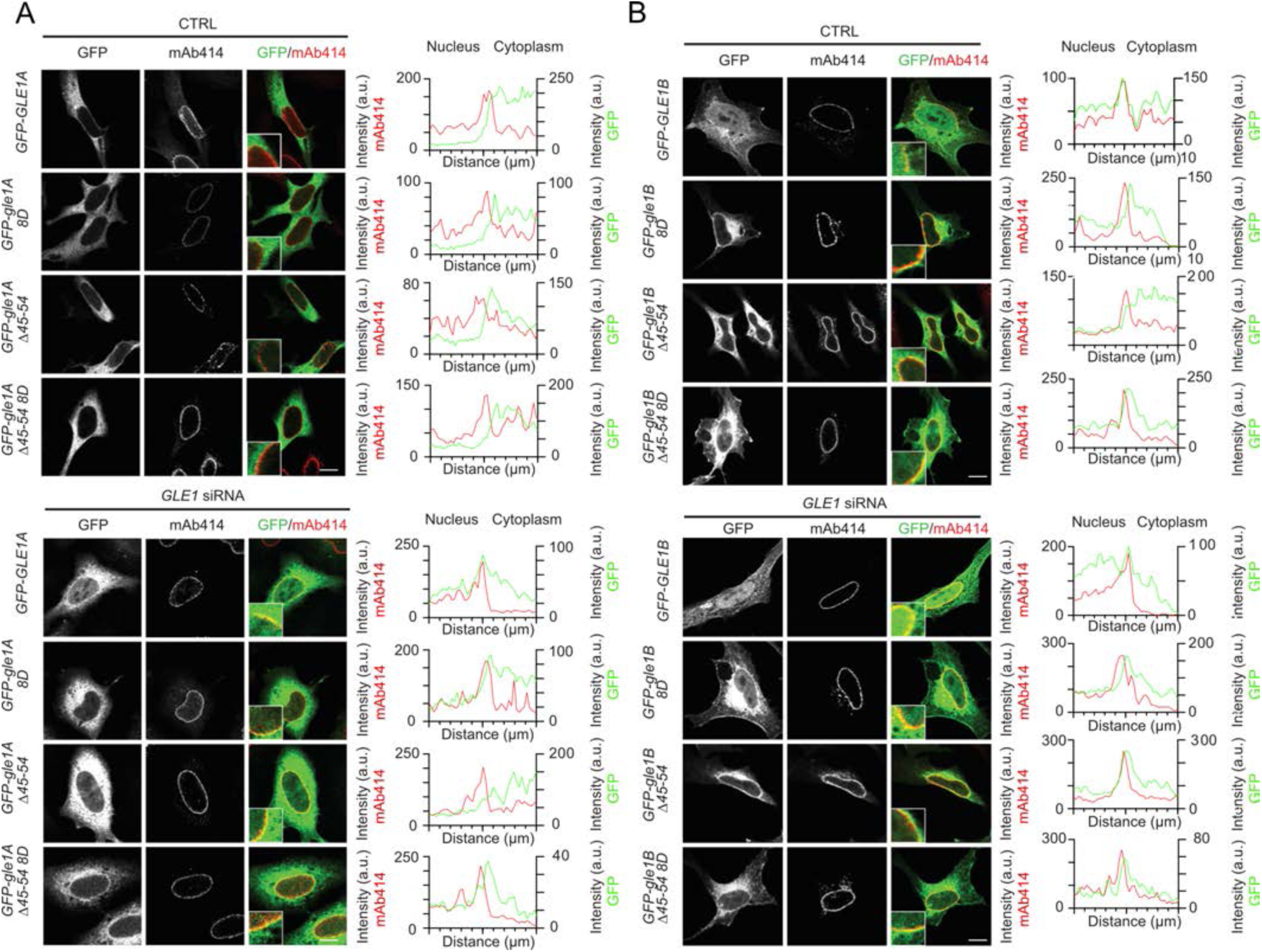
Gle1B oligomerization mutants have altered nuclear rim localization (A, B) CTRL or GLE1 siRNA-treated HeLa cells were transfected with *GFP, GFP-GLE1A, GFP-gle1A-8D, GFP-gle1A-Δ45-54, GFP-gle1A-Δ45-54-8D, GFP-GLE1B, GFP-gle1B-8D, GFP-gle1B-Δ45-54*, or *GFP-gle1B-Δ45-54-8D* plasmids and processed for immunofluorescence using anti-nuclear pore complex proteins antibody (mAb414). Insets highlight nuclear rim with corresponding line intensity plots across the nuclear rim. Scale bar: 10 µm.

As previously reported, in the presence of endogenous Gle1A and Gle1B, exogenously-expressed GFP-Gle1B localized to the nuclear rim, (Fig. 4B) (11). However, by indirect immunofluorescence of GFP with mAB414, we discovered that exogenously-expressed GFP-gle1B-8D, GFP-gle1B-Δ45-54, and GFP-gle1B-Δ45-54-8D only localized near the nuclear rim on the cytoplasmic face but were not colocalized with mAb414 (Fig. 4B). In cells depleted of endogenous Gle1A and Gle1B by *GLE1* siRNA treatment, exogenously-expressed GFP-Gle1B was again colocalized at the NPC with mAb414, whereas GFP-gle1B-8D, GFP-gle1B-Δ45-54, and GFP-gle1B-Δ45-54-8D remained adjacent to the cytoplasmic face (Fig. 4B). This incomplete rim colocalization with the GFP-gle1B oligomerization variants might be related to their loss of steady state intra-nuclear localization. Taken together, these results suggest that proper oligomerization is essential for normal Gle1A and Gle1B steady state localization and association with the NPC.

### Competent mRNA export is dependent on Gle1 oligomerization

Based on our confirmation that impaired oligomerization alters nuclear rim localization of Gle1B, we speculated that defective oligomerization would also perturb mRNA export. Poly(A)^+^ RNA distribution was quantified by *in situ* hybridization with a Cy3-conjugated oligo d(T) in *GLE1* and CTRL siRNA-treated cells, and rescue of the mRNA export defect was analyzed by expressing *GFP, GFP-GLE1B, GFP-gle1B-8D, GFP-gle1B-Δ45-54, or GFP-gle1B-Δ45-54-8D*. Gle1-depleted cells expressing GFP exhibited strong nuclear accumulation of poly(A)^+^RNA, indicative of an mRNA export defect (Fig. 4A). Consistent with previous observation, expression of wild type GFP-Gle1B rescued the export defect (Fig. 4A) (7). However, when the aggregation prone region was deleted from the exogenously expressed protein (GFP-gle1B-Δ45-54) or the coiled-coil domain was disrupted (GFP-gle1B-8D), incomplete rescue of the mRNA export defect was observed (Fig. 5A). When both oligomerization regions were altered (GFP-gle1B-Δ45-54-8D), no rescue of the mRNA export defect was detected, suggesting an additive effect of the two Gle1 oligomerization regions on mRNA export (Fig. 5A, C). These results comprehensively demonstrated that Gle1’s function in mRNA export is profoundly sensitive to its oligomerization state.

**Figure 5.**
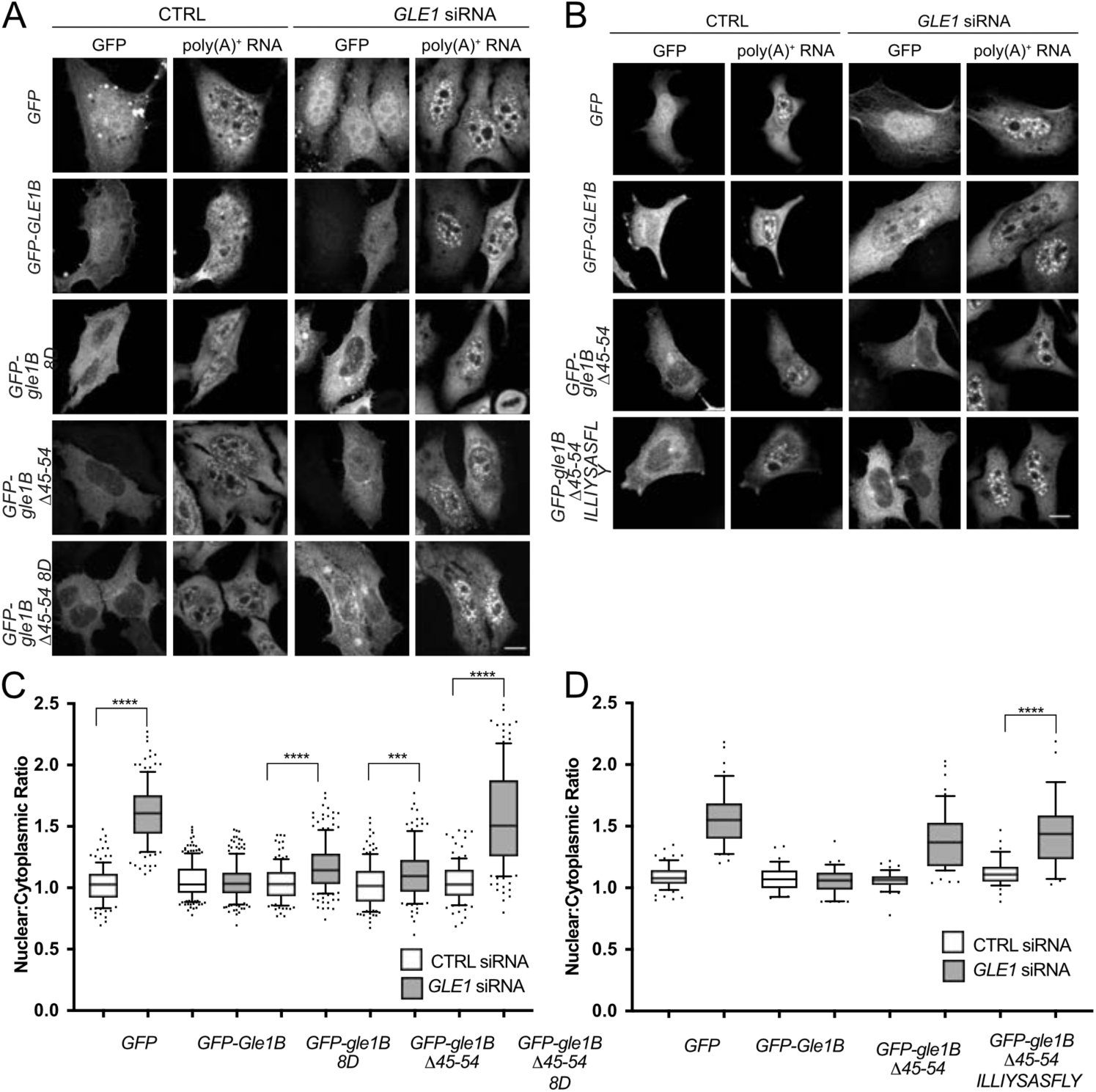
mRNA export is disrupted by Gle1B oligomerization mutants. (A, B) Nuclear poly(A)^+^ RNA accumulation was assessed in CTRL or GLE1 siRNA-treated HeLa cells transfected with *GFP, GFP-GLE1B, GFP-gle1B-8D, GFP-gle1B-Δ45-54, GFP-gle1B-Δ45-54-8D*, or *GFP-gle1B-Δ45-54-ILLIYSASFLY* and detected by in situ Cy3-conjugated oligo-dT hybridization and direct fluorescence microscopy. (C, D) Quantification from three independent experiments of the nulclear:cytoplaspmic ratio of poly(A)^+^ RNA in GFP positive cells and analyzed using the two-tailed, unpaired t-test. ****, p<0.0001; ***; p=0.0004 Error bars represent the standard error of the mean.

To examine the specificity of the aggregation prone domain for Gle1 function, amino acids 45-54 (*LSSYSGQVVE*) were replaced with the unrelated *ILLIYSASFLY* sequence that shows a role in promoting protein aggregation (34). The protein sequence of gle1B-Δ45-54 ILLIYSASFLY was examined with AGGRESCAN and verified that the unrelated sequence in context of Gle1B elicited an aggregation prone hot spot similar to the wild-type sequence (Fig. S1) As with *GFP-gle1B-Δ45-54*, there was an incomplete rescue of the mRNA export defect when *GFP-gle1B Δ45-54 ILLIYSASFLY* was expressed (Fig. 5 B,D). We concluded that the specific sequence of the Gle1 aggregation prone region is required for Gle1 function, and the generic ability to promote aggregation is not sufficient for proper Gle1 function.

### Gle1 oligomerization is required for proper stress granule morphology

Our prior studies show that a phosphorylation cluster in the unstructured N-terminal region of Gle1 influences the function of Gle1 in response to heat-shock stress and also alters the morphology of high molecular weight Gle1 oligomerization discs *in vitro* (16). Because this phosphorylation cluster is positioned between the aggregation prone region and the coiled-coil domain, we investigated whether disrupting Gle1 self-association alters SG characteristics. SGs were induced by heat-shock stress in cells treated with CTRL or *GLE1* siRNA and transiently transfected with *GFP, GFP-GLE1A, GFP-gle1A-8D, GFP-gle1A-Δ45-54*, or *GFP-gle1A-Δ45-54-8D*. SG morphology was visualized by G3BP immunofluorescence, and defects were assessed based on SG number and size. A defect in SG assembly results in an increased number of SGs per cell and decreased size of SGs. As previously shown, a striking SG assembly defect was observed in Gle1-depleted cells, and was rescued by exogenous expression of *GFP-GLE1A* (Fig. 6A) (3). Next, the Gle1A oligomerization mutants were examined for their ability to rescue the SG defects. In CTRL siRNA treated cells, the number and size of heat shock-induced SGs were unaffected (Fig. 6B, C). However, in *GLE1* siRNA treated cells, exogenous expression of the *GFP-gle1A* oligomerization mutants resulted in an increase in SG number. Cells expressing either *GFP-gle1A-8D* or *GFP-gle1A-Δ45-54* showed an incomplete rescue, wherein the number of SGs per cell was significantly increased compared to cells expressing wild type *GFP-GLE1A* but significantly decreased compared to the *GFP* control. Lacking both oligomerization domains, expression of *GFP-gle1A-Δ45-54-8D* resulted in no SG rescue (Fig. 6B). Notably, expressing any one of the three oligomerization mutants in *GLE1* siRNA treated cells gave rise to SGs that are strikingly decreased in average size as compared to *GFP* expressing cells (Fig. 6C). Thus, proper Gle1 oligomerization was crucial for normal stress granule assembly.

**Figure 6.**
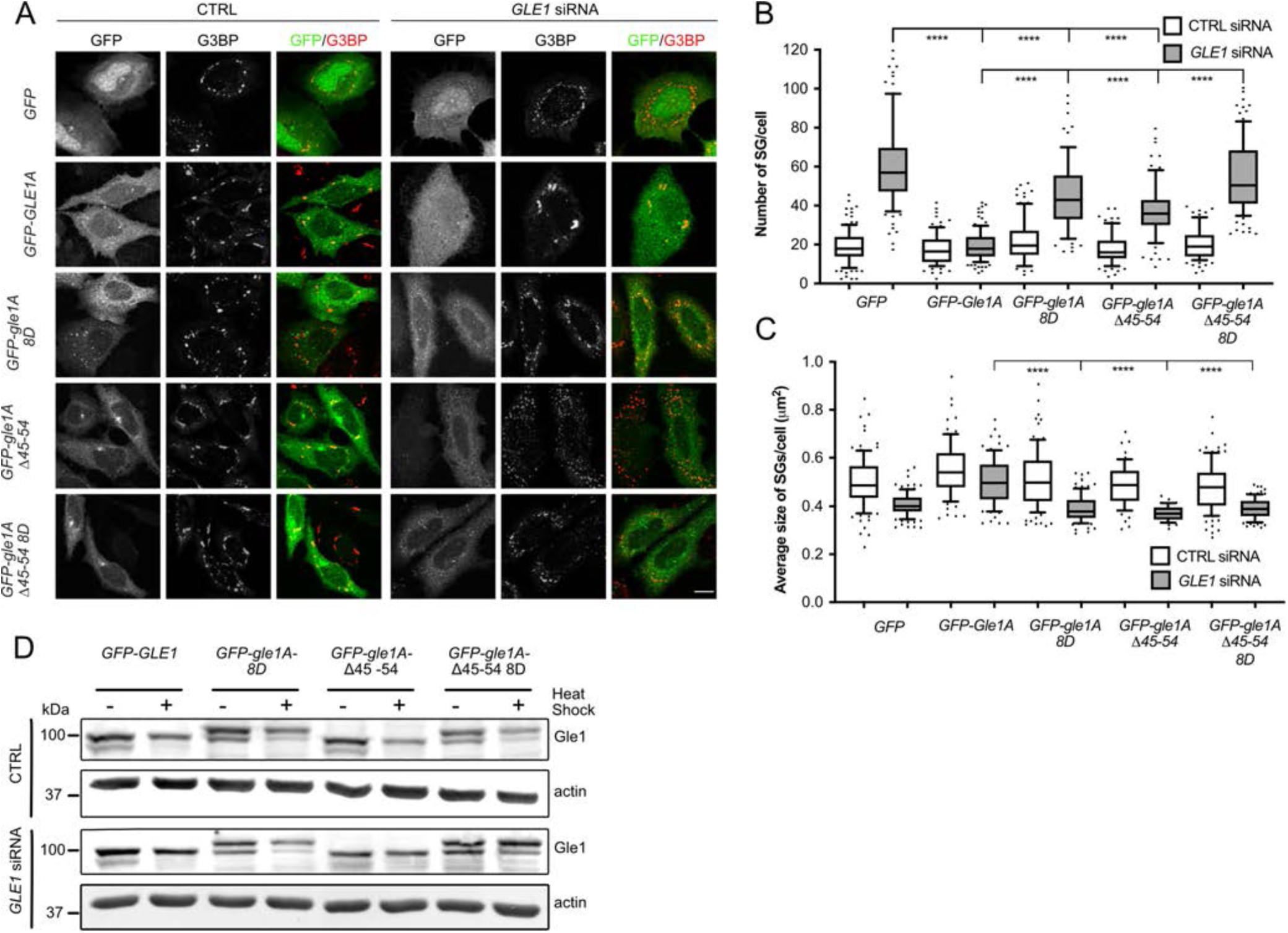
Gle1A oligomerization mutants do not rescue stress granule defects in the absences of endogenous Gle1 but are phosphorylated. (A) HeLa cells treated with either CTRL or GLE1 siRNA were transfected with *GFP, GFP-GLE1A, GFP-gle1A-8D, GFP-gle1A-Δ45-54, or GFP-gle1A-Δ45-54-8D*, subjected to heat shock at 45°C for 60 min, and processed for immunofluorescence using anti-G3BP antibody. Scale bar: 10 µm. Analysis of stress granule number (B) and average size (C). Error bars represents the standard error of the mean from three independent experiments. ****, p<0.0001. (D) Gle1A oligomerization mutants are phosphorylated after heat shock treatment. HeLa cells expressing either *GFP, GFP-GLE1A, GFP-gle1A-8D, GFP-gle1A-Δ45-54*, or *GFP-gle1A-Δ45-54-8D* in CTRL or GLE1 siRNA-treated cells were subjected to heat shock at 45C for 60 or left untreated. Cell lysates were resolved on SDS-PAGE and immunoblotted with anti-Gle1 antibodies.

Given that *gle1* phosphorylation and oligomerization mutants exhibited similar stress granule phenotypes ((16); Fig. 6A), we investigated the phosphorylation status of the oligomerization mutants. Cells treated with either CTRL or *GLE1* siRNA and expressing *GFP, GFP-GLE1A, GFP-gle1A-8D, GFP-gle1A-Δ45-54, or GFP-gle1- Δ45-54-8D* were subjected to heat-shock stress and gle1 phosphorylation levels were monitored by western blot analysis (Fig. 6D). After heat-shock, there was a discernable increase over basal phosphorylation for the respective GFP-Gle1 protein in cells expressing *GFP-GLE1A, GFP-gle1A-8D, GFP-gle1A-Δ45-54, or GFP-gle1A-Δ45-54-8D* that was independent of the presence of endogenous Gle1. Therefore, loss of phosphorylation did not underlie the inability of oligomerization mutants to rescue SG defects.

### Translation activity under stress is regulated by Gle1 oligomerization

Gle1 plays a conserved role in both human and yeast cells in regulating translation initiation (5, 6, 16), and Gle1A specifically regulates translation during the stress response in coordination with SG dynamics (16). Given that Gle1 oligomerization alters SG formation, we speculated that Gle1 oligomerization might also play a role in regulating translation. Overall translation levels in CTRL or *GLE1* siRNA treated cells were measured under nonstress and heat-shock stress conditions by labeling newly translated proteins with L-azidohomoalanine (AHA). Alexa Fluor-488 was covalently attached to AHA using click chemistry as a means of visualization. *mCherry, mCherry-GLE1A, mCherry-gle1A-8D, mCherry-gle1A-Δ45-54, or mCherry-gle1A-Δ45-54-8D* were transiently expressed and translation levels were assessed. In non-stressed cells, the depletion of endogenous Gle1 resulted in a decreased Alexa Fluor-488 signal, indicating a decrease in translation. This defect was rescued by the exogenous expression of *mCherry-GLE1A, mCherry-gle1A-8D, mCherry-gle1A-Δ45-54*, or *mCherry-gle1A-Δ45-54-8D* (Fig. 7A, C). In untreated CTRL siRNA treated cells, no change in translation was observed. As previously reported, when *GLE1* siRNA treated cells were heat-shock stressed, an aberrant increase in translation activity under stress was observed, compared to control siRNA-treated cells (Fig. 7B, D) (Aditi et al., 2015). This defect was rescued by the expression of *mCherry-GLE1A* (Fig. 7B, D). Conversely, expression of *mCherry-gle1A-8D, mCherry-gle1A-Δ45-54* or *mCherry-gle1A-Δ45-54-8D* did not rescue the translation defect, but instead appeared to elevate translation activity (Fig. 7B, D). Taken together, these results suggested that Gle1 oligomerization is not required for its functions in translation under non-stressed conditions; however, proper Gle1 oligomerization is required to regulate translation in response to stress.

**Figure 7.**
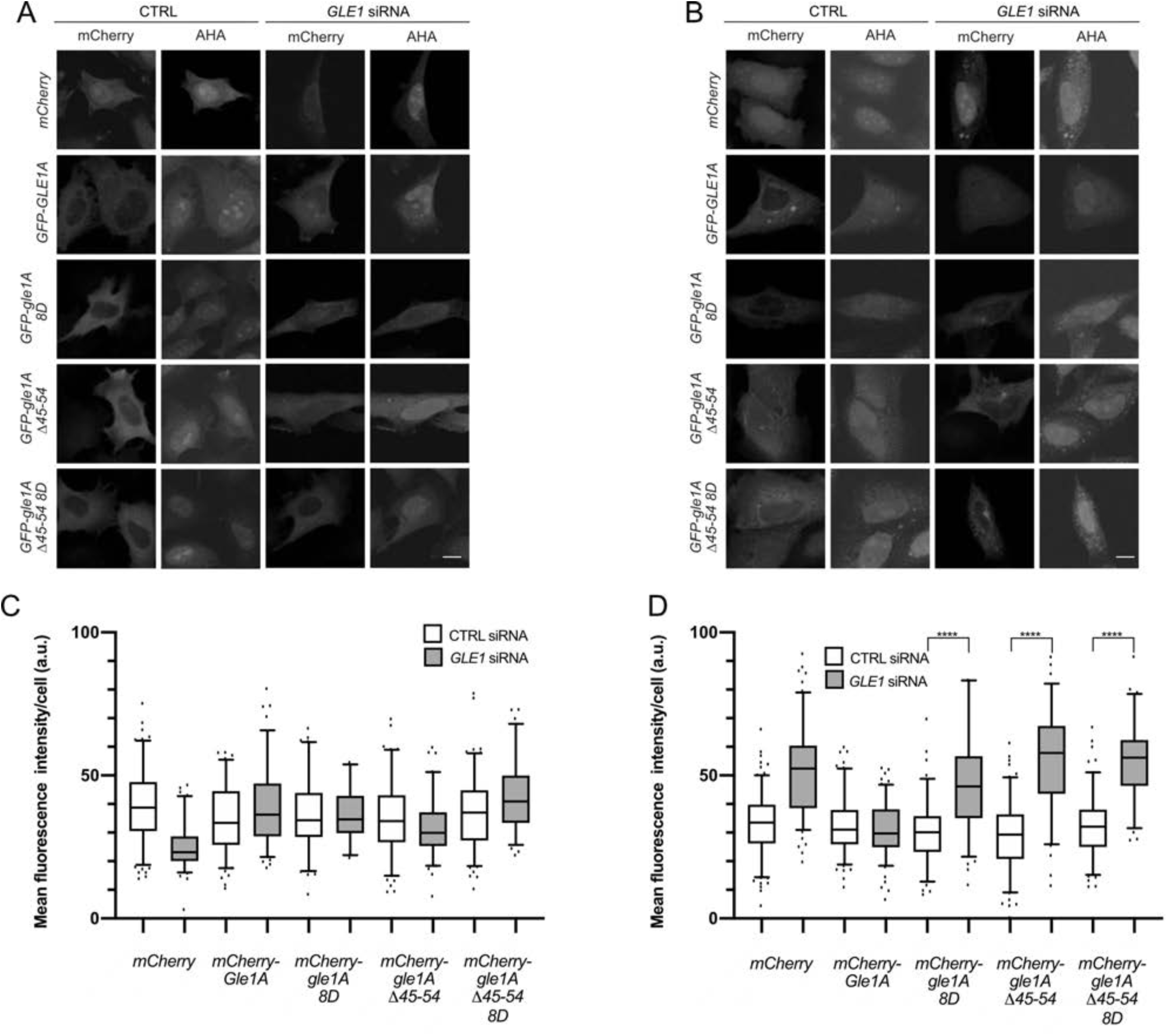
Proper Gle1A oligomerization is required for regulating translation. (A, B) HeLa cells treated with either CTRL or GLE1 siRNA were transfected *with mCherry, mCherry -GLE1A, mCherry -gle1A-8D, mCherry -gle1A-Δ45-54, or mCherry -gle1A-Δ45-54-8D*, subjected to heat shock at 45°C for 60 min (B) or left untreated (A). AHA was added to the incubation for the final 45 min. Samples were processed by Click-iT labeling with Alexa Fluor-488 alkyne for direct fluorescence microscopy. Scale bar: 10 µm. (C, D) Quantification of mean fluorescence intensity of AHA-484 staining in mCherry positive cells. Error bars represents the standard error of the mean from three independent experiments.

## Discussion

In this report, we document the biochemical basis of human Gle1 oligomerization and uncover two distinct regions in the N-terminal region of Gle1 that contribute to self-association through different mechanisms. Both domains are directly required for proper Gle1 function during mRNA export, stress granule assembly, and stress regulated translation. In prior studies, we proposed that the predicted Gle1 coiled-coil domain alone was essential for higher ordered oligomerization (7). Here, we find that while the coiled-coil domain self-associates in a parallel orientation, it is not sufficient for higher ordered Gle1 oligomerization (Fig. 1G). Surprisingly, it is the N-terminal Gle1 IDR upstream of the coiled-coil domain that is necessary and sufficient for producing oligomers of high molecular weight. We identify an aggregation prone patch within amino acids 45-54 of the IDR, and show that deletion of these ten amino acids disrupts higher ordered oligomerization in the context of the entire N-terminal region (Gle1^1-360^) (Fig. 2).

*In vivo*, Gle1 oligomerization through both the aggregation prone region and the coiled-coil domain is essential for proper Gle1 function. Disrupting either region alone or both together results in altered steady state subcellular localization, as shown by a decrease in the nuclear to cytoplasmic ratio and changes at the nuclear rim (Fig. 3,4). These results suggest that oligomerization of Gle1 impacts the nucleocytoplasmic shuttling of Gle1. Oligomerization as a means to regulate nucleocytoplasmic shuttling is observed with other proteins. For example, Ste5 oligomerization promotes nuclear export and function (35) whereas leukemia-associated RhoGEF (LARG) oligomerization prevents nuclear import (36). Interestingly, both of these examples show an increase in cytoplasmic steady state accumulation which is consistent with our data for *gle1* oligomerization-defective mutants. Speculatively, in addition to perturbing Gle1 nucleocytoplasmic shuttling, a decrease in Gle1 homo-oligomerization could also alter interactions with other proteins in the nucleus or cytoplasm and thereby indirectly or directly shift steady state subcellular localization to the cytoplasm.

Proper Gle1B function is essential for efficient mRNA export through NPCs (7). Mutants lacking the aggregation prone region or the coiled-coil domain show a diminished rescue of the mRNA export defect observed when endogenous Gle1 is depleted, whereas disruption of both oligomerization domains results in a complete loss of rescue (Fig. 5). Therefore, the ability of Gle1 to self-associate is required for efficient mRNA export, and the two modes of oligomerization have an additive effect on Gle1 function. During mRNA export, IP_6_-bound Gle1 stimulates DDX19 to promote efficient mRNA export (13). However, *in vitro* studies show that the N-terminal region of Gle1 is dispensable for activating DDX19/Dbp5 ATPase activity (4, 10). As such, we predict that the mRNA export defect observed in *gle1* oligomerization mutants is not due to an inability to activate DDX19/Dbp5. Rather, oligomerization of Gle1 must impact mRNA export in a different manner. Potentially, the altered subcellular localization of Gle1B when oligomerization is impaired might result in mRNA nuclear accumulation by depleting the pool of Gle1 shuttling through the NPC. Alternatively, Gle1 oligomerization might contribute to establishing a DDX19B remodeling platform at the cytoplasmic face of the NPC. A current model based on extensive structural and biochemical data suggests that the mRNA export platform is positioned over the central channel of the NPC at its cytoplasmic face to aid in efficient mRNA export (37), and Gle1 interacts with Nup42 within this complex to position it for stimulating mRNP remodeling (13, 14). Taken together with the results presented here, proper Gle1B oligomerization could be required to position Gle1B over the central NPC channel and/or to create a high local concentration of Gle1B for efficient mRNA export.

In the cytoplasm, Gle1A regulates stress granule dynamics (3, 38). In Gle1-depleted cells, stress granules increase in number and are smaller in size (3). When either the aggregation prone region or the coiled-coil domain is independently disrupted, stress granule assembly defects in Gle1-depleted cells are not fully rescued. Moreover, as was also observed for mRNA export, when there are defects in both oligomerization regions, a complete loss of function is observed (Fig. 5A, B). Interestingly, none of the *gle1A* oligomerization mutants rescue the defect in stress granule size (Fig. 6C). This is consistent with our previous finding that gle1-Fin_major_ is defective in SG assembly (16), and validates our model that Gle1A oligomerization plays an essential role in the fusion step of stress granule assembly. Lastly, Gle1A phosphorylation does not appear to be impacted by impaired oligomerization, as the gle1A oligomerization-defective proteins are phosphorylated under stress conditions in cells (Fig. 5D). Taken with our previous results that SG disassembly is promoted by Gle1A hyperphosphorylation, which in turn destabilizes Gle1A oligomerization (16), our data suggests that both SG assembly and disassembly are highly influenced by the oligomerization state of Gle1A.

Although we previously established that Gle1 regulates translation under non-stressed and stressed conditions (3), it was unclear how the oligomerization state of Gle1 impacts these functions. Here we show that under non-stressed conditions, Gle1 oligomerization is not required for proper translation (Fig. 7A,C). However, under heat shock stress conditions, *gle1A* oligomerization mutants do not properly regulate translation compared to wild type Gle1. Therefore, the distinct roles played by Gle1 in translation are differentially impacted by oligomerization. We speculate that Gle1A oligomerization might impact DDX3 function during translation. In our prior studies (16), we demonstrated that Gle1A phosphorylation alters oligomerization and differentially promotes SG assembly and disassembly potentially through altered regulation of DDX3. In the absence of phosphorylation, Gle1 oligomerizes stably and promotes SG assembly. Upon stress-induced MAPK phosphorylation of Gle1, its self-association is destabilized, DDX3 ATPase activity is inhibited and SGs disassemble. Furthermore, the SG defect observed in Gle1-depleted cells can by rescued by the overexpression of DDX3 (3). These results support a model wherein Gle1A oligomerization is essential to inhibiting DDX3 activity during cellular stress and the coincident global inhibition of translation initiation, and dispensable for DDX3 function during recovery from stress.

Our data also supports a model wherein Gle1A oligomerization is a driving force for proper assembly of stress granules through both coiled-coil interactions and association of the unstructured IDR and aggregation prone regions. Oligomerization of Gle1A might provide a structural scaffold for SG assembly or potentially introduce new protein-protein interactions required for proper SG biology. Although we examined SGs under acute stress conditions, chronic stress conditions elicit stress granules with differing composition and ultimately cell fate (39). Stress granules from acute stress have a pro-survival function (38, 40–42), whereas stress granules from chronic stress have a pro-death function (39). Given that mutations in *GLE1* are linked to neurodevelopmental diseases often resulting in cell death (26, 43), the potential disruption of Gle1A oligomerization would result in SG assembly defects under acute stress, incapacitating the pro-survival effect of stress granules in this scenario. Under chronic stress, which was not examined here, enhanced Gle1 oligomerization could support a pro-death function of SGs by driving the formation of toxic liquid-liquid phase separated granules and perhaps even fibrils. Further studies are needed to address this possibility.

The striking complexity of the Gle1 N-terminal region and its roles in modulating Gle1A and Gle1B function have direct implications for understanding the mechanisms underlying human *gle1* disease-based mutants. The catastrophic effects of the lethal *gle1-Fin*_*major*_ mutation are linked to perturbations of Gle1 oligomerization (7) and, likewise, altered Gle1 oligomerization might also contribute to defects in SG biology and mRNP metabolism that are associated with neurodegenerative disease (44– 47). With two distinct self-association determinants in the N-terminal region, the aggregation prone region and the coiled coil domain, there are multiple mechanisms to control Gle1 oligomerization and impact mRNA export, translation, and SG dynamics through distinct DEAD-box proteins.

## Experimental procedures

### Protein expression and purification

Plasmids (Table S1) were transformed in *Escherichia coli* Rosetta cells under kanamycin and chloramphenicol selection and cultured in Terrific Broth at 37°C until an A600 of 0.8 was reached. Recombinant protein expression was induced with 0.2 mM isopropyl 1thio-β-D-galactopyranoside for 18 hr at 18°C. Cells were harvested by centrifugation, resuspended in 20 mM HEPES (pH 7.5), 500 mM NaCl, 0.5 mM TCEP, and 20% glycerol supplemented with cOmplete EDTA-free protease inhibitor (Sigma-Aldrich, Saint Louis, MO) and 2 mM phenylmethylsulfonyl fluoride, and lysed by sonication. Cleared lysate was loaded on to an amylose resin ((New England Biolabs, Ipswitch, MA) column and washed with 10 CV of lysis buffer. On column cleavage with His-GST-PPS was performed overnight at 4°C. The protease was removed using His-Pur™ Ni-NTA agarose resin (Thermo Fisher Scientific, Waltham, MA). Gle1^152-360^ was further purified by a 30% ammonium sulfate precipitation. The resulting pellet was resuspended in 20 mM HEPES (pH 7.5), 200 mM NaCl, 0.5 mM TCEP, and 10% glycerol and dialyzed overnight at 4°C in the same buffer. Proteins were concentrated and buffer exchanged into SEC buffer (20 mM HEPES (pH 7.5), 200 mM NaCl, 0.5 mM TCEP, and 10% glycerol) prior to S200 size exclusion chromatography. Protein purity was assessed by SDS-PAGE and Coomassie staining throughout the purification process.

MBP-Gle1^1-152^ was eluted from the amylose resin with 10 mM maltose and subsequently buffer exchanged into 20mM HEPES (pH 7.5), 50 mM NaCl, 0.5 mM TCEP, and 10% glycerol. The protein was load onto an ion exchange column (Bio-Rad, Hercules, CA) and eluted with a salt gradient from 50 mM to 1 M NaCl. The peak protein fractions were pooled and buffer exchanged into SEC buffer.

### SEC-MALS

Samples analyzed by SEC-MALS were buffer exchanged by dialysis into SEC-MALS running buffer containing 20 mM HEPES (pH 7.6), 200 mM NaCl, 5% glycerol, 0.5mM TCEP, and 0.05% azide. Samples were applied to a Superose6 10/300 GL (Sigma-Aldrich) column and in-line measurements were recorded for ultraviolet absorbance at 280 nm (using Agilent 1100 series UV detector, Agilent Technologies), static light scattering (DAWN HELEOS 8+light scattering detector, Wyatt Technology), and differential refractive index (Agilent 1200 series refractive index detector, Agilent Technologies) were recorded. System calibration was performed on a BSA standard (Sigma-Aldrich), first resuspended, then buffer exchanged into the SEC-MALS running buffer. SEC-MALS data was analyzed with ASTRA v6.1 software (Wyatt Technology).

### Sequence analysis using web-based servers

The Gle1 amino acid sequence was analyzed for coiled-coil probability and heptad register using MARCOIL and aggregation hotspots using AGGRESCAN as described in (27) and (33), respectively.

### Native PAGE

4-15% acrylamide gradient native gels (8.3 × 7.3 cm) were cast with 3.75mM Tris, pH 8.8. Protein samples were diluted 1:1 with sample solution (0.187M Tris, pH 6.8, 30% glycerol, 80 μg.mL bromophenol blue) and electrophoresed at 150V for 90 min in electrophoresis buffer (0.2M glycine, 0.02M tris). Proteins were visualized by Coomassie staining.

### Circular Dichroism

Circular dichroism experiments were performed with a Jasco J-810 spectropolarimeter outfitted with a Peltier temperature control module. All spectra were collected at 25°C in 20 mM phosphate buffer (pH7.6). Spectra were collected using a 2 nm bandwidth at 1 nm intervals with each data point averaged for 5 s.

### Cell culture manipulations

HeLa cells were grown in complete DMEM (Gibco, Thermo Fisher Scientific) supplemented with 10% fetal bovine serum (FBS, Atlanta Biologicals, Flowery Branch, GA) at 37°C in 5% CO_2_. Knockdown-addback experiments were performed as previously described (7). Briefly, cells were transfected with either a non-targeting siRNA or siRNA targeting *GLE1* using HiPerFect (Qiagen, Germantown, MA). Expression plasmids (Table S1) were transiently transfected using Fugene6 (Promega, Madison, WI) according to the manufacture’s recommendations. For stress granule analysis, cells were treated at 45°C for 60-min in a non-CO_2_ air incubator as previously described. Translation was assessed by replacing the media with DMEM absent of cysteine and methionine and incubated for 45 min at 37°C. Cells were then left untreated or subjected to heat-shock for 15 min prior to the addition of the methionine analogue L-azidohomoalane (AHA, 50 μM) (Thermo Fisher Scientific) and incubations continued for an additional 30 min.

### Live cell microscopy

Cells used for live cell imaging were plated in 35 mm No 1.5 glass bottom dishes from MatTek prior to transient transfection. Phenol red-free DMEM (Gibco) supplemented with 10% FBS was applied, and imaging was performed using a Zeiss LSM 710 META inverted microscope with a plan-apochromat 63x/1.4 oil-immersion objective (Zeiss, White Plains, NY).

### In situ hybridization, immunofluorescence, and AHA labeling

Cells were plated on No 1.5 round coverslips in a 24-well plate prior to knockdown-addback. Following a 72 hr post-siRNA treatment with a final 24 hr transient transfection, cells were fixed with paraformaldehyde and permeabilized with 0.2% Triton X-100. Poly(A)^+^ RNA was localized by Cy3-conjugated oligo d(T) hybridization as previously described (7). For immunofluorescence, permeabilized cells were blocked with 10% FBS/PBS for 1 hr at room temperature. G3BP was detected with anti-G3BP (1:300, BD Biosceince, San Jose, CA) and the nuclear rim was detected with mAb414 (1:300, Biolegend, San Diego, CA). Alexa Fluor 594-conjugated secondary antibodies (Thermo Fisher Scientific) were used at 1:1000. Images were acquired on a Leica TCS SP5 confocal microscope using a 63x/1.4 NA oil-immersion objective and a digital charge coupled device camera. AHA-labeled proteins were detected with 1 μM alkyne-tagged Alexa Fluor-488 using Click-iT cell detection reagents (Thermo Fisher Scientific) for 45 min at room temperature as previously described (3).

### Quantification of microscopy

All images were processed with ImageJ (National Institutes of Health, Bethesda, MA). Poly(A)^+^ localization was quantified in GFP fluorescent positive cells by the average Cy3-conjugated oligo d(T) intensity in the nucleus and cytoplasm. Stress granule number and size were determined using the ImageJ 3D objects counter plug-in as previously described (3). AHA-labeled protein levels were quantified by determining the average cellular intensity of Alex Fluor-488.

### Immunoblotting

HeLa cells were plated in 60 mm dishes and knockdown-addback were performed as above with indicated siRNA and plasmids. For phosphorylation studies, cells were incubated at 45°C for 60 min. Cells were washed in TBS and lysed with RIPA buffer. Cleared lysates were loaded onto 7% gels for SDS-PAGE. Following electrophoresis, proteins were transferred to PVDF and probed with anti-Gle1 (ASW47.1) and detected with goat anti-rabbit IRDye 680RD (LI-COR Biosciences, Lincoln, NE).

## Data availability

All data are contained within the article.

## Acknowledgements

We thank members of the S. R. W. and Yi Ren (Vanderbilt University) laboratories for thoughtful discussions and advice.

## Funding and additional information

This work was supported by the National Institutes of Health Grant 5R37GM051219 (to S. R. W.). Microscopy was performed in part through the use of the Vanderbilt Cell Imaging Shared Resource, which is supported by the National Institutes of Health grants CA68485, DK20593, DK58404, DK59637, and EY08126. The content is solely the responsibility of the authors and does not necessarily represent the official views of the National Institutes of Health.

## Abbreviations and nomenclature

IP_6_: inositol hexakisphosphate
mRNA: messenger RNA
mRNP: messenger ribonucleaoprotein
Dbp: *Saccharomyces cerevisiae* DEAD-box protein
DDX: human DEAD-box protein
NPC: nuclear pore complex
SG: stress granule
IDR: intrinsically disordered region
LCCS1: lethal congenital contracture syndrome 1
SEC-MALS: size exclusion chromatography multiangle light scattering
SDS-PAGE: sodium dodecyl sulfate polyacrylamide gel electrophoresis
SEC: size exclusion chromatography
EM: electron microscopy
MBP: maltose-binding protein
GFP: green fluorescent protein
CTRL: control
MAPK: mitogen-activated protein kinase
TCEP: tris(2-carboxyethyl)phosphine
Ni-NTA: nickel-nitrilotriacetic acid
HEPES: 4-(2-hydroxyethyl)-1-piperazineethanesulfonic acid
AHA: L-azidohomoalaine
FBS: fetal bovine serum
PBS: phosphate-buffered saline
TBS: tris-buffered saline
RIPA: radioimmunoprecipitation assay buffer
PVDF: polyvinylidene fluoride

